# Eco-evolutionary dynamics of prior selfing rate promotes the coexistence without niche partitioning under reproductive interference

**DOI:** 10.1101/2020.04.07.029082

**Authors:** Koki R. Katsuhara, Yuuya Tachiki, Ryosuke Iritani, Atushi Ushimaru

## Abstract

1. When the two or more plants species share the same pollinators, pollinator-mediated reproductive interference make coexistence difficult. Recent studies suggested prior autonomous selfing mitigate reproductive interference, could enabling coexistence without pollination niche partitioning (pre-emptive selfing hypothesis). However, there are no studies to test whether evolution of prior selfing promote the coexistence, considering eco-evolutionary dynamics of population size, selfing rate and inbreeding depression.
2. To examine conditions that the evolution of prior selfing promote coexistence under mutual reproductive interference especially in the point of view for pollinator availability and dynamics of inbreeding depression, we constructed individual-based model in which two plant species compete against each other in the form of mutual reproductive interference and can evolve prior autonomous selfing rate. We expected that purging of deleterious mutations could cause evolutionary rescue because inferior species could rescue population density through the evolution of prior selfing if the strength of inbreeding depression decreases with an increase of population’s selfing rate.
3. Our simulation demonstrated that the evolution of prior selfing could promote the coexistence while reproductive interference caused competitive exclusion without evolution. We found that lower pollinator availability tended to prefer rapid evolutionary shift to higher prior selfing rate, it neutralizes the negative effect of reproductive interference, and population dynamics exhibit neutral random walk in both species. When the strength of inbreeding depression decreased with an increase in population’s selfing rate, moderate pollinator availability resulted in long-term coexistence in which relative-abundance-dependent selection on the prior selfing rate rescue population density of inferior species intermittently.
4. *Synthesis*. We showed that the evolution of prior selfing could increase population growth rate of inferior species and consequently enable the long-term coexistence with evolutionary rescue. This is the new mechanisms explaining co-evolutionary coexistence of closely related plant species without niche partitioning and consistent with recent studies reported that closely related mixed-mating species are sympatrically growing even under the mutual reproductive interference.

## Introduction

Clarifying the conditions under which competing species can coexist is a traditional and most fundamental subject in ecology (May, 1974; Chesson, 2000). Numerous empirical and theoretical works have shown that niche partitioning between competing species is required for their coexistence: i.e. the intraspecific competition should be larger than interspecific competition (Chesson, 2000; Silvertown, 2004). Closely related species, which potentially share the same or very similar resources and reproductive habits, are therefore expected to heavily compete against each other, likely being unable to coexist (Gröning & Hochkirch, 2008; Burns & Straus, 2011; Whitton, Sears & Maddison; 2017).

In flowering plants, when two or more plant species depend on the same pollinators for reproduction, interspecific pollinator-mediated pollen transfer can cause strong reproductive interference that results in competitive exclusion of either species or niche partitioning between species (Levin & Anderson, 1970; Takakura, Nishida, Matsumoto & Nishida, 2008; Runquist & Stanton, 2013; Moreira-Hernández & Muchhala, 2019). Reproductive interference via interspecific pollen transfer can occur when both or either of increase in heterospecific pollen deposition on the stigma and conspecific pollen loss on heterospecific flower (Mitchell, Flanagan, Brown, Waser & Karron, 2009; Morales & Traveset, 2008). Specifically, Pollen deposition from closely related heterospecies is known to strongly decrease female reproductive success owing to pollen tube growth competition in the style, ovule discounting and/or hybridization (Harder, Cruzan & Thomson, 1993; Nishida, Kanaoka, Hashimoto, Takakura & Nishida, 2014; Whitton et al., 2017). Thus, reproductive interference via heterospecific pollen deposition may favour spatiotemporal segregation in flowering or floral trait displacement, with a consequence that they use different, or different body parts of the same, pollinator species (e.g. Runquist 2012; van der Niet & Johnson 2012; Huang & Shi, 2013).

Selfing has gathered much recent attention as an alternative mechanism that can mitigate reproductive interference by heterospecific pollen transfer from competing relatives (Fishman & Wyatt, 1999; de Waal, Anderson & Ellis, 2015; Katsuhara & Ushimaru, 2019). Recent studies have suggested that selfing constitutes a reproductive barrier among sympatrically coexisting related species (Fishman & Wyatt, 1999; Martin & Willis, 2007; Goodwillie & Ness 2013; Brys, van Cauwenberghe & Jaquemyn, 2016). Selfers with small and inconspicuous flower (selfing syndrome) which therefore receives fewer pollinator visits are likely to coexist with outcrossing relatives (Sicard & Lenhard, 2011; Kalisz et al., 2012). Thus, reduced heterospecific pollen deposition owing to fewer visits and/or reproductive assurance via self-pollination might mitigate the negative effect of reproductive interference in selfers, although it is difficult to clarify their relative importance in general (Fishman & Wyatt, 1999; Martin & Willis, 2007; de Waal et al., 2015; Brys et al., 2016).

Recent studies further hypothesize that “prior” rather than “delayed” autonomous selfing can mitigate the negative effect of reproductive interference via interspecific pollen transfer and promote species coexistence independent of the presence of pollinator visitations (the pre-emptive selfing hypothesis; Randle, Spigler & Kalisz, 2018; Katsuhara & Ushimaru, 2019). Theoretical and empirical studies have suggested that prior selfing unlikely evolves with frequent pollinator visitations (Lloyd, 1992; Kalisz, Vogler & Hanley, 2004; Eckert et al., 2010). However, in the presence of reproductive interference by an abundant competitor species, frequent pollinator visitations largely reduce outcrossing success of an inferior species. In such a situation, the evolution of prior selfing can mitigate the negative effect of reproductive interference from the competitor and would rescue the inferior species from competitive exclusion.

The pre-emptive selfing hypothesis should be tested in the context of eco-evolutionary dynamics of population size, selfing rate and inbreeding depression. Because the negative effect of reproductive interference that decreases outcrossing success becomes greater with an increase in the relative abundance of competing species (Levin & Anderson, 1970; Katsuhara & Ushimaru, 2019), population dynamics of mutually competing species should be an important driving factor of the evolution of prior autonomous selfing of a given species. The evolution of prior selfing could rescue the population density of competitively inferior species via mitigation of reproductive interference while it could also result in self-extinction due to the negative effect of inbreeding depression on population growth rate depends on the strength of inbreeding depression (Cheptou, 2019; Katsuhara & Ushimaru, 2019). Therefore, dynamics of inbreeding depression is an important factor influencing the evolution of selfing because the strength of inbreeding depression is often expressed as a decreasing function of population’s sefling rate due to “purging” of deleterious, recessive alleles (Schemske & Lande, 1985; Chaelesworth, Chaelesworth & Morgan, 1990; Lloyd, 1992; Husband & Schemske, 1996; Crnokrak & Barrett, 2002; Goodwillie, Kalisz & Eckert, 2005; Charlesworth & Willis, 2009). Thus, dynamics of population size, the degree of selfing rate and inbreeding depression of competing species are ideally considered to examine the adaptive significance of prior selfing under reproductive interferences. To the best of our knowledge, however, no studies have examined the eco-evolutionary dynamics of these variables, and therefore little is known about the possibility of coexistence under reproductive interference, followed by evolution of prior selfing.

In this study, to examine the pre-emptive selfing hypothesis, we constructed a model in which two plant species sharing the same pollination niche and can evolve prior autonomous selfing compete against each other in the form of mutual reproductive interference (i.e. eco-evolutionary dynamic model). Using the model, we addressed following questions. Can prior selfing evolve under mutual reproductive interference and promote their coexistence as an evolutionary rescue agent? Is inbreeding depression an important determinant for the joint dynamics of population size and selfing rate? By answering to these questions, we discuss the conditions in which the evolution of prior selfing promotes the long-term coexistence of closely related species sharing the same pollination niche.

## Model

### Community structure, pollination, seed production and germination processes

We develop an individual-based model of competition between two annual flowering plant species (species with discrete generation) within a site whose carrying capacity is *K*: *K* individuals of both or either of species 1 (sp_1_) and 2 (sp_2_) lives in the site for each generation (default *K* value is 2,000). Here, the relative abundance of sp_*i*_ at the *t*-th generation are denoted as *q*_*i,t*_ where *i* is either 1 or 2 and *q*_1,*t*_ + *q*_2,*t*_ = 1 holds. Thus, the number of individuals of sp_*i*_ equals *K q*_*i,t*_. In the model, we assume that ecological niches of sp_1_ and sp_2_ completely overlap with each other, although the two species produce no hybrids.

First, we describe the pollination and fertilization processes in the model. Each individual of both species produces *n* ovules which are fertilized via prior autonomous selfing and outcrossing mediated by pollinators and *g* pollen grains. The *j*-th individual of sp_*i*_ fertilizes their ovules via prior autonomous selfing at the rate of *r*_*i,j,t*_ in the *t*-th generation (1 ≤ *j* ≤ *Kq*_*i,t*_ for *i* = 1 or 2). Thus, an integral number of ovules obtained by rounding *nr*_*i,j,t*_ are fertilized via prior selfing and the others were remained for pollinator-mediated outcrossing. Here, we assumed that an integral number obtained by rounding proportion *P* (0 ≤ *P* ≤ 1) of the *n*(1 – *r*_*i,j,t*_) ovules are pollinated with outcrossed conspecific and/or heterospecific pollen grains by pollinators. Using *P* < 1, we can formulate pollinator limitation. Here, we assume that pollinators indiscriminately visit flowers of both species and carried their pollen in proportion to their relative flower abundances.

Pollen parent of each outcrossed ovule of individual *j* is randomly assigned to conspecies with the probability of 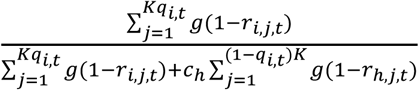, where *c*_*h*_, *r*_*i,j,t*_ and *r*_*h,j,t*_ (0 ≤ *c*_*h*_, *q*_*i,t*_, *r*_*i,j,t*_ and *r*_*h,j,t*_ ≤ 1) are the strength of reproductive interference from heterospecies (sp_*h*_) and the rates of prior selfing of the *j*-th individuals of sp_*i*_ and sp_*h*_, respectively. The parameter *c*_*h*_ is interpreted as the competitive ability of a heterospecific pollen grain relative to that of a conspecific one to get fertilization with individual *j*’s ovule. Besides, the probabilities of which pollen grains of individual *j* fertilize conspecific and heterospecific ovules are described as 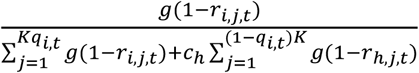 and 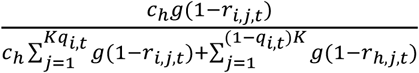, respectively. In this formulation, we assume that outcross pollen proportionally decreases with the prior selfing rate in respective individuals of both species. This assumption mimics increased pollen discounting and/or anther-stigma interference with increasing the prior selfing rate (Lloyd and Webb, 1986; Webb and Lloyd, 1986; Karron, Jackson, Thumser & Schlicht 1997; Fishman, 2000; Barrett 2002). Because we assumed random pollination, it should be noted the ovule of *j*-th individual is fertilized by pollen of *j*-th individual (pollinator-mediated self-pollination; geitonogamy) with the above probability and they are treated as self-fertilized pollen as well as the ovules fertilized by prior autonomous selfing.

Second, we denote the seed production process. We assume that only ovules fertilized by self- and outcrossed-conspecific pollen can develop seeds whereas those fertilized by heterospecific pollen produce no seeds. A cost of selfing relative to outcrossing is also assumed as follows. In sp_*i*_, all outcrossed ovules develop sound seeds whereas selfed ovules set seeds at the rate of 1 – *I*_*i,t*_, where *I*_*i,t*_ (0 ≤ *I*_*i,t*_ ≤ 1) is the strength of inbreeding depression at generation *t* in sp_*i*_. Here, *I*_*i,t*_ is described as a function of the population’s selfing rate at the (*t* – 1)th generation of sp_*i*_, *S*_*i,t*–1_:

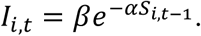

*S*_*i,t*–1_ is calculated as total number of selfed sound seeds divided by total number of sound seeds in sp_*i*_ in the last generation. *I*_*i,t*_ can be interpreted as an evolutionary variable, which decreases with an increase in the population’s selfing rate in the parental generation owing to the accumulation–purging balance of deleterious mutations (Schemske & Lande, 1985; Husband & Schemske, 1996; Crnokrak & Barrett, 2002). The coefficient *α* expresses how inbreeding depression steeply decreased with increasing the population’s selfing rate whereas the intercept *β* (0 ≤ *β* ≤ 1) indicates the level of inbreeding depression when complete outcrossing occurs in the population. We simulated various *α* and *β* values to test various scenarios in the eco-evolutionary dynamics of population size and selfing rate.

As the final process, *K* seedlings from all seeds produced by both species are randomly selected and construct generation *t* + 1. We assume no interspecific differences in competitive ability at germination and establishment processes as well as other ecological and genetic features: *c*_*h*_, *α*, and *β* are also equal for both sp_1_ and sp_2_. In addition, our model has no spatial structure.

### Inheritance and mutation of the rate of prior autonomous selfing

To describe the evolutionary dynamics of prior autonomous selfing, our model assumes that the prior selfing rate of individual *j* in the next generation *r*_*i,j,t*+1_ is determined as the parental average. Thus, the prior selfing rate is assumed to be a quantitative genetic trait value which can be influenced by various quantitative traits such as the degrees of herkogamy and/or dichogamy and the proportion of cleistogamous flowers (Culley & Klooster, 2007; Kalisz et al., 2012). In addition, *r*_*i,j,t*+1_ can be mutated to be slightly lower or higher than the parental mean (a random value between -*s* and +*s* is added to the parental mean) with a probability *µ. µ* and *σ* are the rate and effect size of mutation, respectively. We used 0.05 and 0.1 for *µ* and *σ* as default values, respectively. If mutated *r*_*i,j,t*+1_ becomes larger than 1 or smaller than 0, we use the values 1 and 0, respectively.

### Simulation settings and categorization of eco-evolutionary consequence

To explore conditions for the coexistence of the two species, we examined the effects of pollinator availability (*P*) and inbreeding depression-selfing rate relationship (*α* and *β*) on the consequences of evolution. We tested two following scenarios for inbreeding depression-selfing rate relationship. *I*_*i,t*_ is fixed (*α* = 0; *β* = 0.1, 0.3, 0.5, 0.7 or 0.9) or it varies in concert with the population’s selfing rate (*α* = 0, 0.5, 1, 2, 4 or 8; *β* = 0.9), with the whole parameter range of 0 ≤ *P* ≤ 1 (Table 1). In each simulation run, the initial numbers of individuals for both species are equal as *K*/2. The initial autonomous selfing rates for individuals were generated randomly with normal distribution whose mean and standard deviation are *r*_intial_ (1/2) and *sd*_intial_ (1/6) for both species. Each run continues for 2,000 generations or until either species goes extinct.

**Table 1.** List of parameters.

| Parameter | Definition | Default value |
| --- | --- | --- |
| $r_{i,j,t}$ | The ratio of ovules fertilized via prior autonomous selfing in the $j$-th individual of $sp_i$ at the $t$-th generation. | 0-1 |
| $q_{i,t}$ | Relative abundance of $sp_i$ at the $t$-th generation | 0-1 |
| $P$ | Pollinator availability | 0-1 |
| $c_h$ | Strength of reproductive interference | 1 |
| $\alpha$ | Slope of inbreeding depression function | 0, 0.5, 1, 2, 4, 8 |
| $\beta$ | Intercept of inbreeding depression function | 0.1, 0.3, 0.5, 0.7, 0.9 |
| $\mu$ | mutation rate | 0.05 |
| $\sigma$ | Effect size of mutation | 0.1 |
| $K$ | Carrying capacity (number of individual plant) | 2000 |
| $n$ | Number of ovules per individual plant | 200 |
| $r_{\text{initial}}$ | Mean of initial prior autonomous selfing rate | 0.5 |
| $sd_{\text{initial}}$ | Standard deviation of initial prior autonomous selfing rate | 1/6 |

After 50 simulation runs for each parameter setting, we classified the eco-evolutionary dynamics into four categories based on ecological and evolutionary status of the species. When the simulation terminated by the extinction of either species and the population mean of prior autonomous selfing rate in survivors was higher or lower than 0.5, the result was categorized as (1) competitive exclusion by selfer or (2) that by outcrosser, respectively. Meanwhile, the runs in which the two species still coexisted at the 2,000th generation were also divided into two categories, (3) a coexistence with evolutionary rescue by prior selfing and (4) a coexistence with neutral dynamics, based on following procedures.

To detect the evolutionary rescue, we calculated the population growth rate and selection gradient in each generation of sp_*i*_. Population growth rate (*W*_*i,t*_) for the *t*-th generation is calculated as *Kq*_*i,t*+1_/*Kq*_*i,t*_ (= *q*_*i,t*+1_/*q*_*i,t*_). For clarifying the selection gradient on the prior selfing rate, we identified seed and pollen parents of all seeds and calculate a correlation coefficient between selfing rate *r*_*i,j,t*_ and seeding and siring success of each individual as the selection gradient (*G*_*i,t*_) at the *t*-th generation. The positive (or negative) *G*_*i,t*_ means that the higher (or lower) rate was adaptive at the generation in sp_*i*_. Then, the evolutionary rescue by prior selfing is defined as a state following two conditions are satisfied simultaneously: (1) a significant negative correlation between *q*_*i,t*_ and *G*_*i,t*_ (i.e., a population decline facilitates the evolution of selfing), (2) a significant positive correlation between the population mean of prior selfing rate 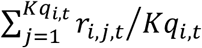 and *W*_*i,t*_ (i.e., the evolution of selfing increases population growth). Significances of these two correlations were examined by a permutation test. We permutated the variables of the last 500 generations in each run 10,000 times and made null distribution and the 95% prediction interval of each correlation to test the significance of the value of the run. When both or either of the correlation coefficients were not significant, the run was categorized into the coexistence with neutral dynamics (Fig. 1).

**Fig. 1.**
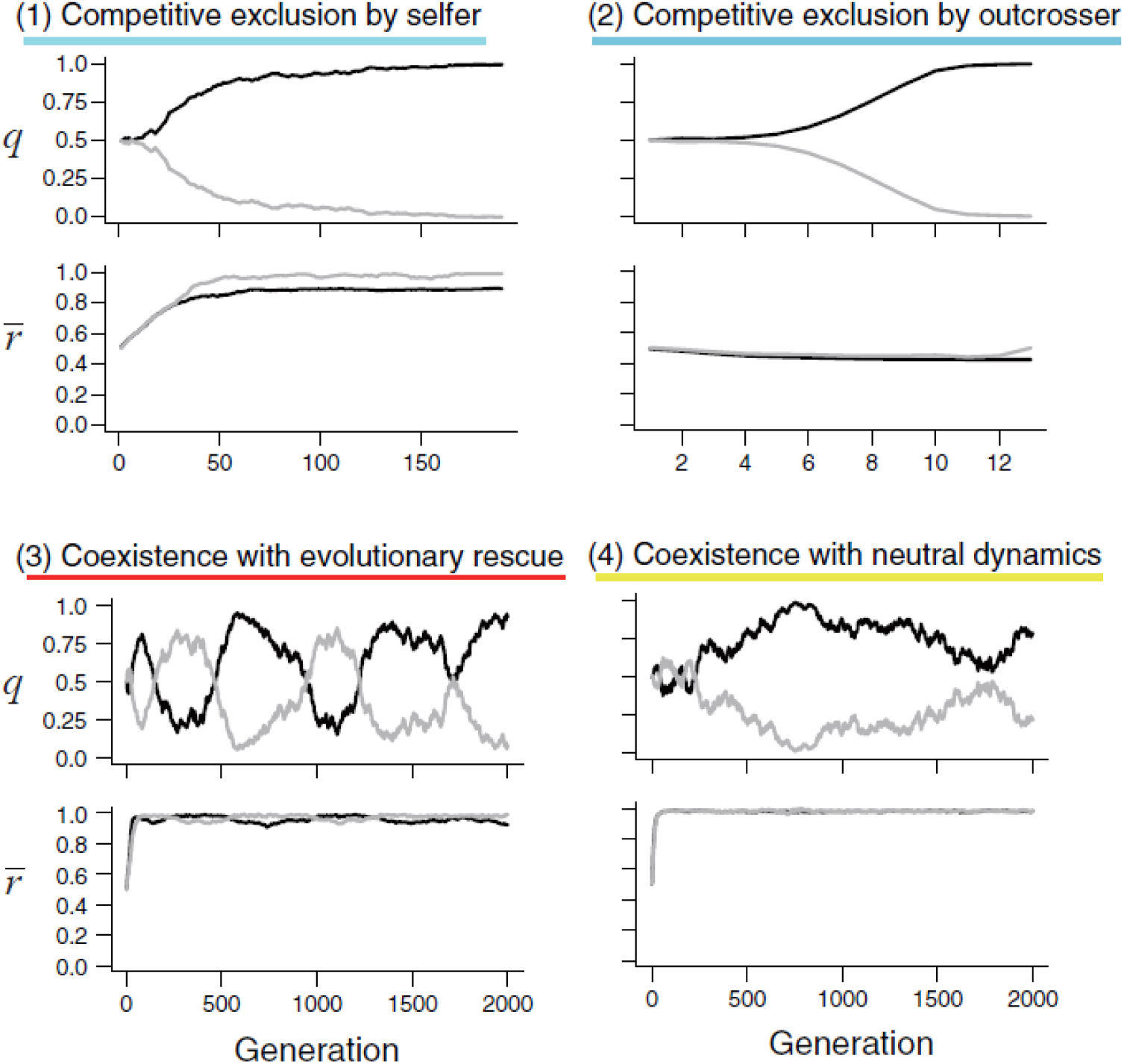
Examples of four consequences of eco-evolutionary dynamics in our simulations: (1) competitive exclusion by selfer; (2) competitive exclusion by outcrosser; (3) coexistence with evolutionary rescue; (4) coexistence with neutral dynamics. Upper and lower graphs of each category show relative abundance and population mean of prior selfing rate dynamics of sp_1_ (black line) and sp_2_ (grey line), respectively.

We compared difference in long-term stability between coexistences with neutral dynamics and evolutionary rescue. We selected a typical parameter set for each coexistence type: *P* = 0.4, *α* = 0.5, *β* = 0.9 for that with evolutionary rescue (ER set) and *P* = 0.1, *α* = 4, *β* = 0.9 for coexistence with neutral dynamics (ND set). For each parameter set, we conducted 200 simulations for 10,000 generations and recorded the generation until which two species coexisted.

We also checked how simulation results change depending on the strength of reproductive interference and the initial population’s mean selfing rate. We examined simulations in which *c*_h_ (= 0.0, 0.25, 0.5, 0.75 or 1.0) and *r*_initial_ (= 0.0, 0.25, 0.5, 0.75 or 1.0) varied with the above parameter settings (ER and ND sets) and run 50 simulations for each parameter set. Moreover, to check the population dynamics of the two species with the fixed population’s prior selfing rates, we conducted simulation runs in which sp_1_ and sp_2_ had the same or different fixed prior selfing rates (0 ≤ *r*_initial_ ≤1) with the same two parameter settings (ER and ND sets). Finally, we run simulations with the fixed abundance of two species to examine the effect of fixed population size on the evolution of prior selfing rate in the two parameter settings (ER and ND sets).

## Results

### Eco-evolutionary dynamics with fixed inbreeding depression

We found that lower pollinator availability preferred the evolution of higher selfing rate in both species, often promoting their coexistence with neutral dynamics (Fig. 2). Conditions for the coexistence with neutral dynamics was more limited by higher inbreeding depression (Fig. 2). The coexistences with neutral dynamics were always realized when the two species evolved the prior selfing rate close to 1.0, which neutralized their mutual reproductive interference (Fig. 1). During the coexistence, population dynamics of both species exhibited a random walk. Therefore, the coexistence with neutral dynamics is not stable in the long term and the extinction of either species occurred when simulations continued for more generations (see the section below, *Long-term stability of the coexistences*). Meanwhile, when either or both of *I* and *P* are large, the eco-evolutionary dynamics tended to be terminated by competitive exclusion (Fig. 2). Especially when both of *I* and *P* are large, competitive exclusion by outcrosser always terminated the eco-evolutionary dynamics (Fig. 2). The coexistence with evolutionary rescue rarely occurred when the inbreeding depression was fixed and independent of the population’s selfing rate (Fig. 2).

**Fig. 2.**
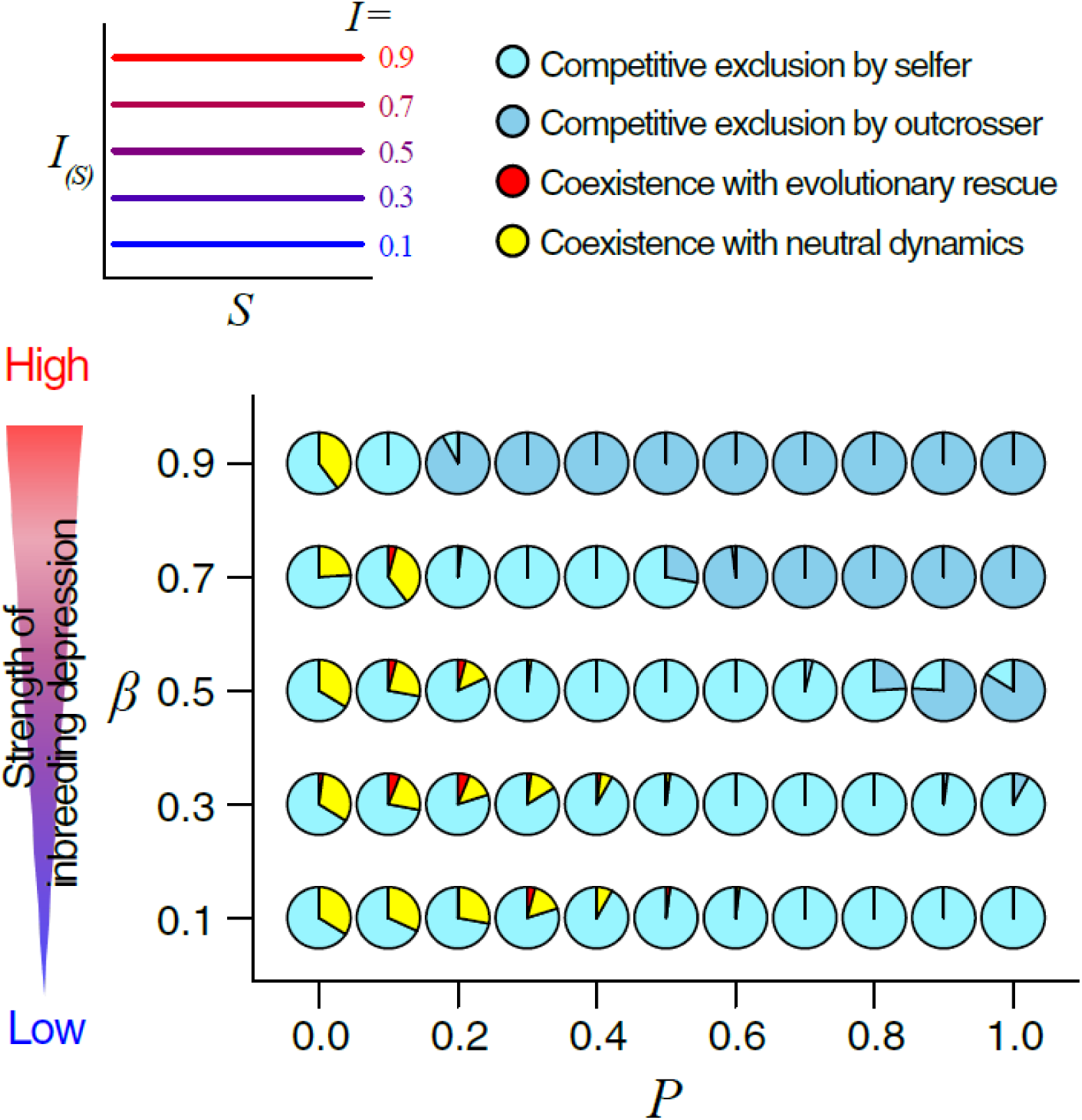
Effects of pollinator availability and the strength of inbreeding depression on simulation consequence in fixed inbreeding depression scenarios. Each pie chat shows frequencies of four categories of eco-evolutionary consequences of 50 simulation runs (Fig. 1).

### Eco-evolutionary dynamics with variable inbreeding depression

In the scenarios with variable inbreeding depression, conditions for both types of coexistence were more relaxed compared to those assuming fixed inbreeding depression (Figs. 2, 3). Interestingly, conditions with intermediate levels of pollinator availability and the slope of inbreeding depression function *α* more frequently facilitated the coexistence with evolutionary rescue or neutral dynamics than other conditions (Fig. 3).

**Fig. 3.**
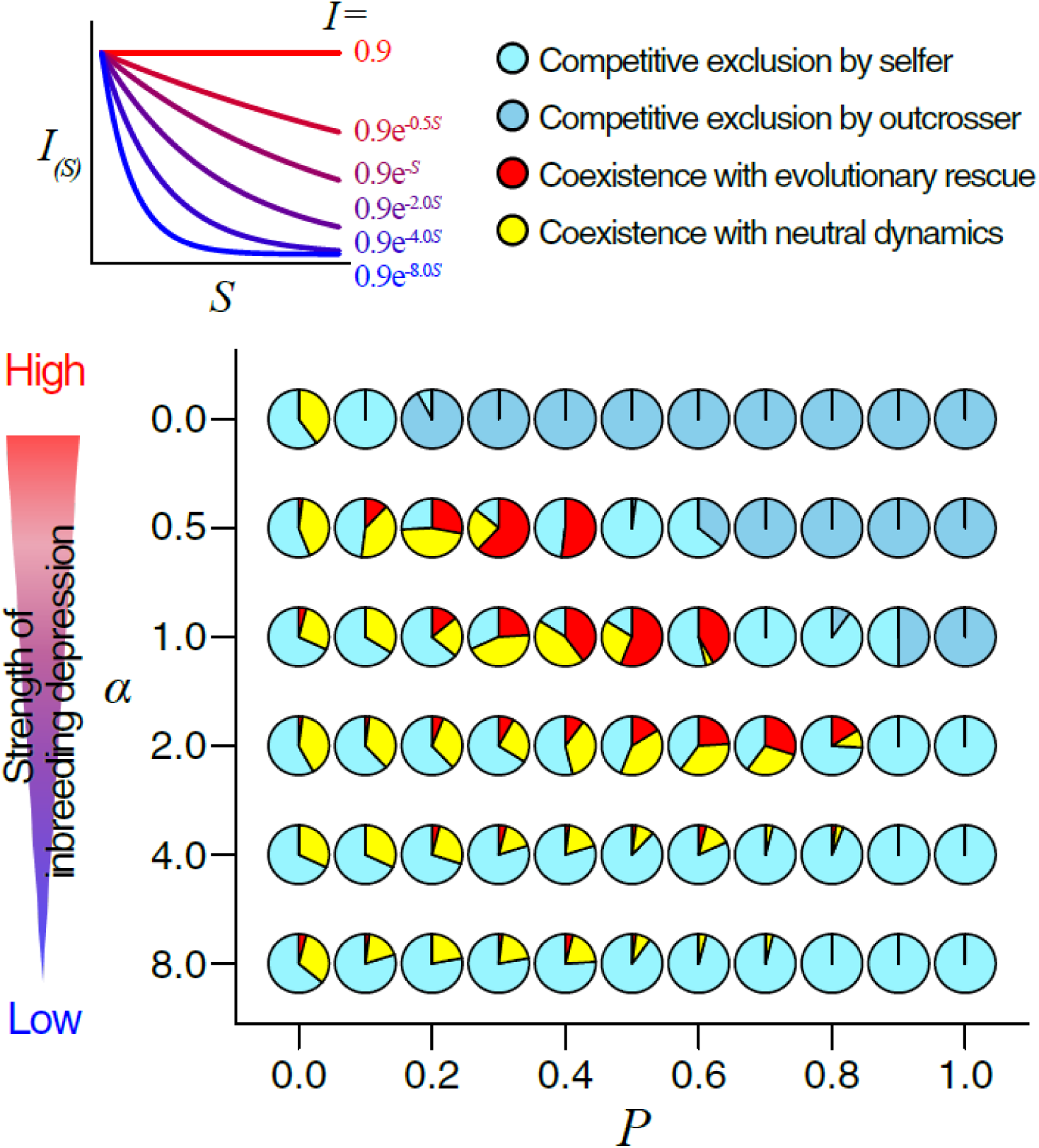
Effects of pollinator availability and the strength of inbreeding depression on the simulation consequence in variable inbreeding depression scenarios. Each pie chat shows frequencies of four categories of eco-evolutionary consequences of 50 simulation runs (Fig. 1).

When inbreeding depression sharply decreases as the population’s selfing rate increases (*α* ≥ 4.0), competitive exclusion by selfer occurred in wider conditions as in those with lower fixed inbreeding depression (*α* = 0 and *β* = 0.1 or 0.3; Figs. 2, 3). Meanwhile, when inbreeding depression more gently decreased with increasing the population’s selfing rate (*α* = 0.5), competitive exclusion by outcrosser tended to occur in the presence of higher pollinator availability like in the cases both of *I* and *P* are large in fixed inbreeding depression scenario.

### Long-term stability of the coexistences with neutral dynamics and evolutionary rescue

The coexistence with evolutionary rescue continued until the 10,000-th generation if the fluctuations of the relative abundances (*q*_*i,t*_) and the prior selfing rates (*r*_*i,t*_) have once started, while the coexistence with neutral dynamics never coexisted before reaching the 10,000-th generation (Fig. 4).

**Fig. 4.**
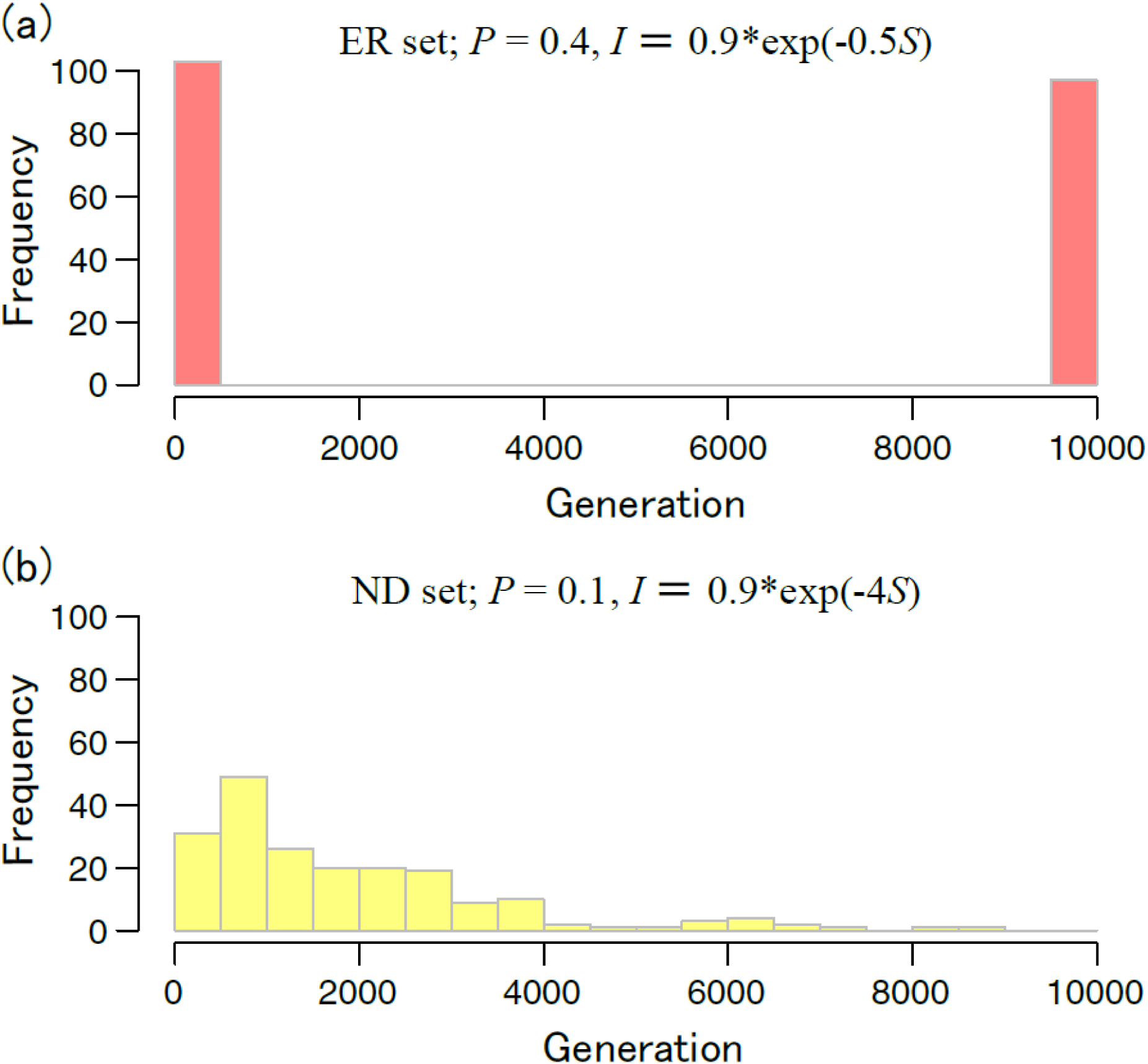
Histograms of generation until when two species coexisted in two typical parameter sets for coexistences with evolutionary rescue (a, ER set) and with neutral dynamics (b, ND set), respectively. The graphs made from the outcomes of 200 simulation runs.

### *Dependence of simulation consequences on c*_h_ *and r*_*initial*_

In the simulations with ER sets, we found that coexistence with evolutionary rescue nearly always occurred with high initial population’s selfing rate (*r*_initial_ ≥ 0.75) and presence of mutual reproductive interference (*c*_h_ > 0.0). When the initial population’s selfing rate was low (*r*_initial_ ≤ 0.25), the both types of coexistence rarely or very infrequently occurred in both the ER and ND sets (Fig. 5). Moreover, no competitive exclusion by outcrosser was found when the initial population’s selfing rate was high (*r*_intial_ ≥ 0.75). Meanwhile, the strength of reproductive interference (*c*_h_) seems unlikely to largely influence the coexistence with neutral dynamics with the ND parameter setting. However, the coexistence of evolutionary rescue never occurred without mutual reproductive interference (*c*_h_ = 0.0) with the ER set.

**Fig. 5.**
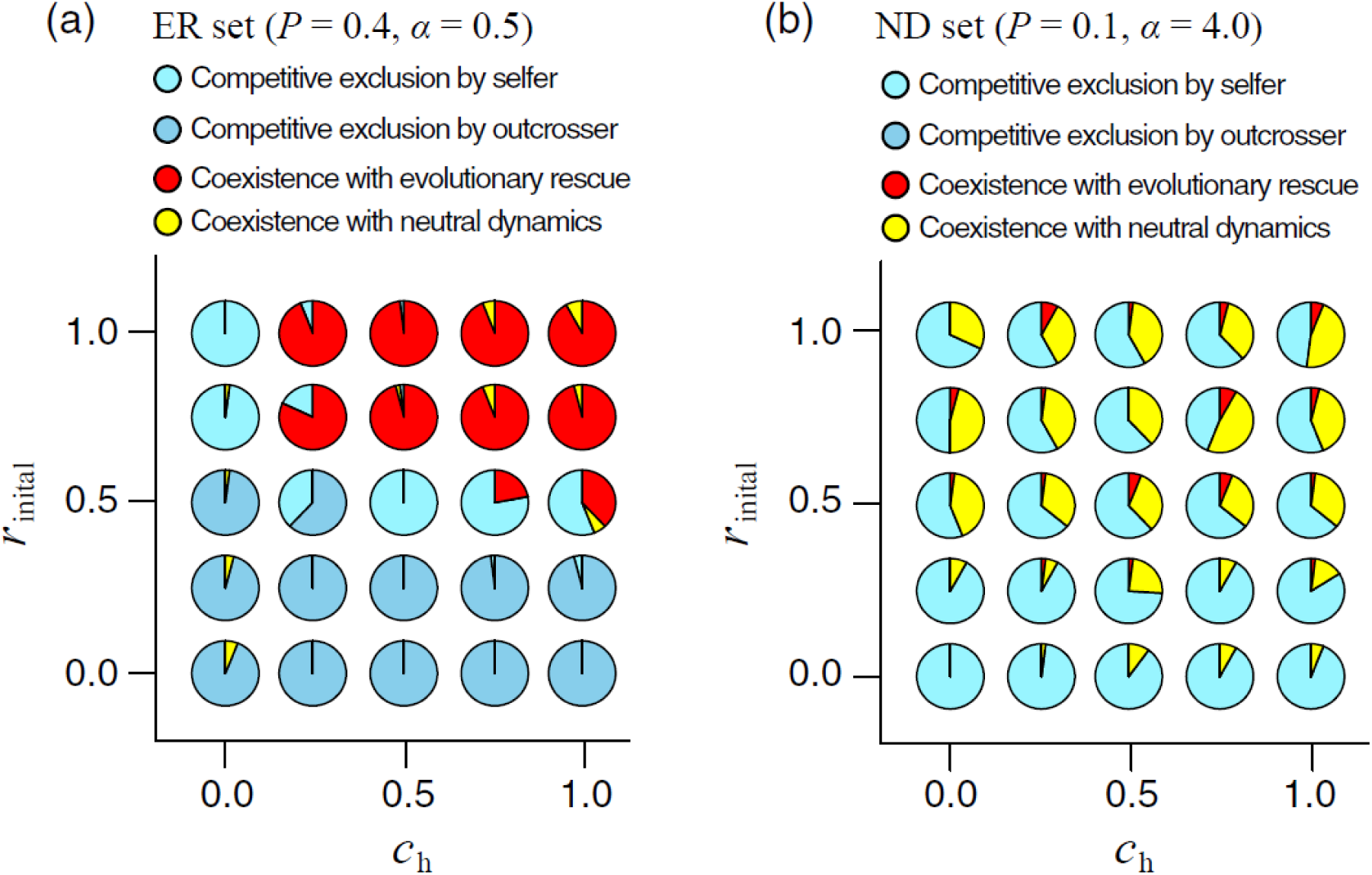
Effects of strength of reproductive interference and initial population’s mean selfing rate in two typical parameter sets for coexistences with evolutionary rescue (a, ER set) and with neutral dynamics (b, ND set), respectively. Each pie chat shows frequencies of four categories of eco-evolutionary consequences of 50 simulation runs (Fig. 1).

### Simulation consequences with fixed prior selfing rate and fixed population size

We found that coexistence for 2,000 generations very rarely occurred when the prior selfing rates were fixed in the two species for the both parameter settings except when both species had the same and very high prior selfing rates (Fig. 6). Winners were always the species having higher prior selfing rates with the ND parameter set whereas winners were usually the species having the lower and higher prior selfing rates in the below and above areas of the line of *r*_2_ = - *r*_1_ + 0.6, respectively, with the ER set (Fig. 6). In the simulations with the fixed abundance of two species, the evolutionary shift to the higher prior selfing rate was favored only when the relative abundance of focal species was lower than 1/2 with the ER parameter set (Fig. 7). Meanwhile, under the ND set, very high prior selfing rate was always favored independent on their abundance (Fig. 7).

**Fig. 6.**
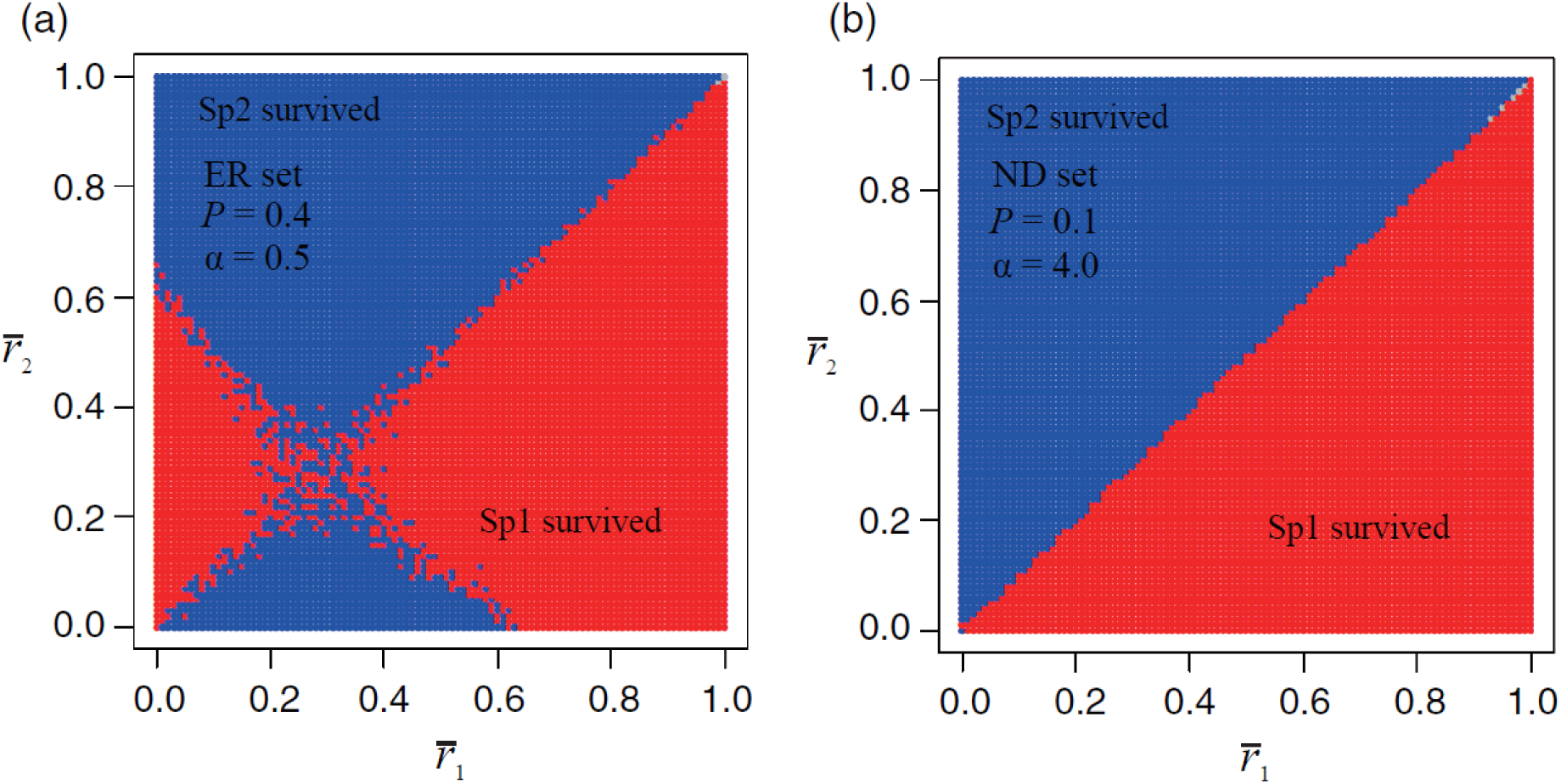
Ecological consequences with fixed population’s prior selfing rate in two typical parameter sets for coexistences with evolutionary rescue (a, ER set) and with neutral dynamics (b, ND set), respectively. X and Y axes indicate population’s mean prior selfing rate of sp_1_ and sp_2_, respectively. Blue and red areas mean that survivor is sp_1_ and sp_2_, respectively, and grey regions (shown upper right corner of each panel) indicated that coexistence continued for 2,000th generations.

**Fig. 7.**
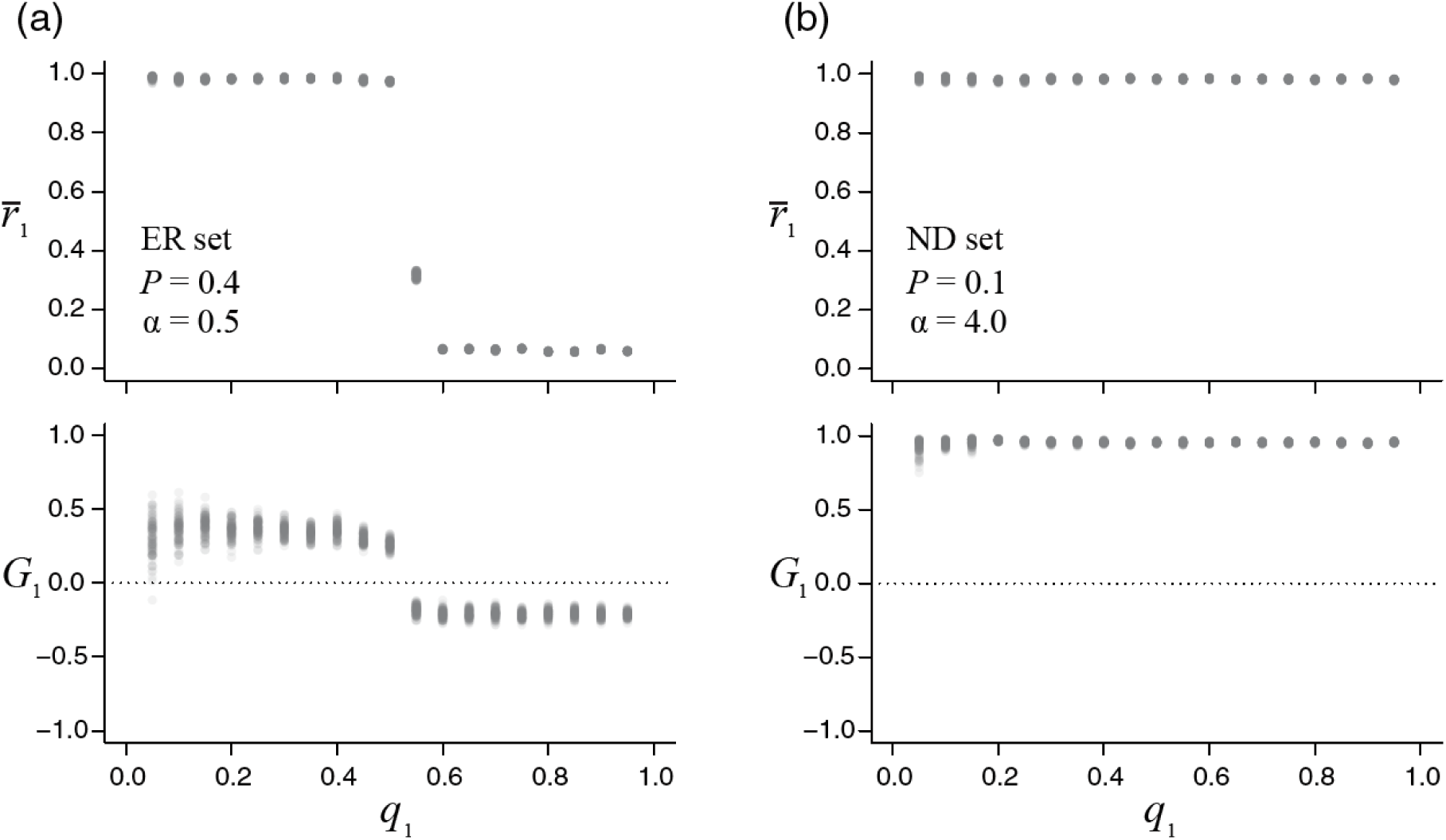
Evolutionary consequences with fixed relative abundance in two typical parameter sets for coexistences with evolutionary rescue (a, ER set) and with neutral dynamics (b, ND set), respectively. Upper and lower graphs indicate the correlations between population’s mean prior selfing rate and selection gradient, and relative abundance of the focal species in last 500 of 2,000 generations, respectively.

## Discussion

Our model revealed that the evolution of prior selfing can promote the coexistence in the presence of mutual reproductive interference while the coexistence rarely occurred without the evolution of prior selfing (Figs. 2, 3, 6). In the variable inbreeding depression scenario (inbreeding depression decreases with an increase in the population’s selfing rate), both types of coexistence tended to be more occurred than in the fixed inbreeding depression scenario when comparing same pollinator availability (Figs. 2, 3). Especially when the strength of inbreeding depression gently decreased and pollinator availability was intermediate level, the coexistence with evolutionary rescue often occurred and stably continued for very long-term (Figs. 3, 4).

Firstly, we discuss the processes enabling the coexistence with evolutionary rescue in our model (Fig. 1). At the early generations, the stochastic process makes slight difference in population size and selfing rate between the two species and reproductive interference by more abundant species with higher selfing rate enlarge the difference. In such a situation, low outcross success due to increased heterospecific pollen deposition facilitates the evolution of high prior selfing rate in the inferior species. This evolutionary shift toward high selfing rate improves the population growth rate via an increase in selfed seed production (i.e., evolutionary rescue by prior selfing occurs) especially when inbreeding depression is weakened with increasing the population’s selfing rate in the inferior species. Meanwhile, once becoming the dominant species, reproductive interference from the competitor is getting weaker so that inbreeding depression favors lower prior selfing rate in the species. Lower prior selfing rate, in turn, can reduce total seed production of the population when pollinator availability is not high, leading to lower population growth rate compared to the competitor. This relative-abundance-dependent selection on the prior selfing rate promotes a negative relationship between and fluctuations of the prior selfing rate and population size through generations. Due to this out-of-phase fluctuations occurring both in two species, the long-term coexistence of the two species is realized under mutual reproductive interference.

Here, suitable conditions for the coexistence with evolutionary rescue are discussed by comparing to empirical knowledge. Our simulation demonstrated that the coexistence with evolutionary rescue occurred with moderate pollinator limitation, variable and moderate levels of inbreeding depression, the relatively higher initial prior selfing rate and the presence of reproductive interference. High pollinator availability always favors competitive exclusion by either outcorsser or selfers depending on the level of inbreeding depression. In other words, the long-term coexistence under the reproductive interference requires pollinator limited conditions which are prevailing in wild flowering plants (Larson & Barrett, 2000). Gently variable inbreeding depression still function as the cost of selfing even when the population’s selfing rate of given species is very high. While inbreeding depression due to deleterious recessive alleles are thought to be rapidly purged with increasing population’s selfing rate, weak late acting inbreeding depression caused by weakly deleterious mutations and heterozygous advantage due to overdominance cannot be purged even in predominantly selfing species (Charlesworth et al., 1990; Husband & Schemske, 1996; Crnokrak & Barrett, 2002; Charlesworth & Wills, 2009). Additionally, although it may not be surprising, we found that higher initial prior selfing rate widens the possibility of the coexistence with evolutionary rescue (Fig. 5). The finding suggests that only a pair of predominantly selfing or of mixed-mating species can coexist stably under reproductive interference, being consistent with recent studies on the coexistence under mutual reproductive interference (Tokuda et al., 2015; Katsuhara & Ushimaru, 2019; Nishida et al. unpublished data). Without reproductive interference, this type of coexistence never occurred even when other parameter settings are suitable for the coexistence (Fig. 5). This result is very interesting and proposes that mutual reproductive interference can act the cost of outcrossing and promoting more selfing (Katsuhara & Ushimaru, 2019), likely making a fluctuation pattern in the prior selfing rate throughout the generation.

The coexistence with neutral dynamics was often found in conditions with lower pollinator availability and weak fixed or moderately variable inbreeding depression (Figs. 2, 3). In such conditions, the higher prior selfing rate evolves very quickly to be almost completely 1.0 in both species (Fig. 1), which should be free from the negative effect of reproductive interference from competitor. Both species exhibit population dynamics of neutral random walk (Hubbell, 2001; Chave, 2004) and coexist, so that stochastic events will stop this type of coexistence at some point in time (Fig. 4). In our model, this type of coexistence was usually found in the parameter conditions where competitive exclusion by selfer frequently occurred, suggesting that these consequence categories do not differ qualitatively (Figs. 2, 3, 5). The rate of evolutionary change in prior selfing rate differed between these categories and the coexistence occurred when the high prior selfing rate evolved more rapidly in both species (Fig. 1). Many predominately selfing weeds usually coexist in human-disturbed habitats where pollinators are often limited (Baker, 1974), most likely being explained by this type of coexistence. Empirical studies have shown the evolutionary shift to higher prior selfing rate (often via reduction of herkogamy) can rapidly occurred under pollinator limitation (Roels & Kelly, 2011; Brys & Jacquemyn, 2012; Gravasi & Schiestl, 2017; Cheptou, 2019). To apply our results to selfing-species coexistence in the field, the rate of evolutionary change of the prior selfing rate and pollinator availability are better to be examined in future studies.

Under conditions with high pollinator availability and strong inbreeding depression, mutual reproductive interference causes very rapid competitive exclusion by outcrosser, being consistent with expectations in the previous works that considered no limitation in outcross gamete transfer (Fig. 1, 2, 3; Levin & Anderson, 1970; Kishi & Nakazawa, 2013). Besides, competitive exclusion by selfer is frequently occurred under conditions with weak inbreeding depression and/or low pollinator availability. The exclusion occurred more slowly comparing to the exclusion by outcrosser (Fig. 1). The difference was likely due to that reproductive interference no more reduced seed production in highly selfing species.

In both types of coexistence, co-evolutionary shifts to extremely high prior selfing rate (over 0.9) was necessary in both competing species (Fig. 1). Many previous empirical studies, however, reported coexistences of an extremely selfer and a related outcrosser (Fishman & Wyatt, 1999; Brys et al., 2016; Randle et al., 2018). This difference between the field observations and our results might be explained by in two possible mechanisms which are not assumed in our model. First, some kinds of ecological differences, such as competitive ability for germination and strength of inbreeding depression, might exist between the study species, promoting the coexistence of species with different mating systems. Second, selfers in these studies always exhibited reduced floral attractiveness which minimized the negative effect of reproductive interference via receiving few pollinator visits (Fishman & Wyatt, 1999; Brys et al., 2016; Randle et al., 2018). Meanwhile, our model assumed only the rate of prior selfing was evolvable as in the previous model (Cheptou, 2019). Such an evolution of selfing floral syndrome could enable obligative selfers to coexist with outcrossing relatives. Some recent studies reported the mutual reproductive interference between two sympatrically growing mixed-mating species which have showy flowers with frequent pollinator visitations and traits promoting prior autonomous self-pollination (cleistogamous flowers in *Impatiens noli-tangare* and *I. textori*, Tokuda et al., 2015; bud pollination in *Commleina communis* and *C. c*. f. *ciliata*, Katsuhara & Ushimaru, 2019). The coexistences found in these study systems could be explained by prior-selfing mediated evolutionary rescue, which are predicted by our model. To test this possibility, monitoring of eco-evolutionary dynamics of these competing species in the fields will be required. Although the complete test will require much time and effort, to examine the relationships among population’s selfing rate, inbreeding depression and relative abundance in the field should improve our understanding of co-evolutionary coexistence mechanisms without pollination niche partitioning as the first step.

In conclusion, our model successfully showed that the evolution of prior selfing could increase population growth rate of inferior species and consequently enable the long-term coexistence with evolutionary rescue. We successfully showed that evolutionarily variable inbreeding depression based on accumulation–purging balance of deleterious mutations expand the possibility of coexistence and promote the long-term coexistence. The result suggests that genetic dynamics of inbreeding depression within a given species may largely influence dynamics of community where pollinator-mediated competition occurs. Finally, we propose new mechanisms explaining co-evolutionary coexistence of closely related species under mutual without any kinds of niche differentiation and spatial structures. The applicability and generality of the proposed mechanisms should be investigated empirically in future.

## Acknowledgements

This study was performed with support from a Grant-in-Aid for JSPS Fellows to K.R.K., JSPS KAKENHI (No. 16H04845, 17K15197, & 20K15876) and JSPS Overseas Research Fellowship to Y.T., JSPS KAKENHI (No. 19K22457 & 19K23768) to R.I. and JSPS KAKENHI (No. 16K07517 & 19K06855) to A. U. We are grateful to the following people for their very helpful comments: N. Ohmido, Y. Takami, T. Minamoto, S. Sugiura, M. K. Hiraiwa, H. S. Ishii and T. Y. Ida.

## Reference

Barrett, S. C. H. (2002). Sexual interference of the floral kind. Heredity, 88, 154–159. doi: 10.1038/sj.hdy.6800020

Baker, H. G. (1974). The Evolution of Weeds. Annual Review of Ecology and Systematics, 5(1), 1–24. doi: 10.1146/annurev.es.05.110174.000245

Brys, R., & Jacquemyn, H. (2012). Effects of human-mediated pollinator impoverishment on floral traits and mating patterns in a short-lived herb: an experimental approach. Functional Ecology, 26, 189–197. doi: 10.1111/j.1365-2435.2011.01923.x

Brys, R., van Cauwenberghe, J., & Jacquemyn, H. (2016). The importance of autonomous selfing in preventing hybridization in three closely related plant species. Journal of Ecology, 104, 601–610. doi: 10.1111/1365-2745.12524

Burns, J. H., & Strauss, S. Y. (2011). More closely related species are more ecologically similar in an experimental test. Proceedings of the National Academy of Sciences, 108, 5302–5307. doi: 10.1073/pnas.1013003108

Charlesworth, B., Charlesworth, D., & Morgan, M. T. (1990). Genetic loads and estimates of mutation rates in highly inbred plant populations. Nature, 347, 380–382. doi: 10.1038/347380a0

Charlesworth, D., & Willis, J. H. (2009). The genetics of inbreeding depression. Nature Reviews Genetics, 10, 783–796. doi: 10.1038/nrg2664

Chave, J. (2004). Neutral theory and community ecology. Ecology Letters, 7, 241–253. doi: 10.1111/j.1461-0248.2003.00566.x

Cheptou, P.-O. (2019). Does the evolution of self-fertilization rescue populations or increase the risk of extinction? Annals of Botany, 123, 337–345. doi:10.1093/aob/mcy144

Chesson, P. (2000). Mechanisms of Maintenance of Species Diversity. Annual Review of Ecology and Systematics, 31, 343–366.

Crnokrak, P., & Barrett, S. C. H. (2002). Perspective: Purging the Genetic Load: A Review of the Experimental Evidence. Evolution, 56, 2347–2358. doi: 10.1111/j.0014-3820.2002.tb00160.x

Culley, T. M., & Klooster, M. R. (2007). The cleistogamous breeding system: A review of its frequency, evolution, and ecology in angiosperms. The Botanical Review, 73, 1. doi: 10.1663/0006-8101(2007)73[1:TCBSAR]2.0.CO;2

de Waal, C., Anderson, B., & Ellis, A. G. (2015). Relative density and dispersion pattern of two southern African Asteraceae affect fecundity through heterospecific interference and mate availability, not pollinator visitation rate. Journal of Ecology, 103, 513–525. doi: 10.1111/1365-2745.12358

Eckert, C. G., Kalisz, S., Geber, M. A., Sargent, R., Elle, E., Cheptou, P.-O., … Winn, A. A. (2010). Plant mating systems in a changing world. Trends in Ecology & Evolution, 25, 35–43. doi: 10.1016/j.tree.2009.06.013

Fishman, L. (2000). Pollen Discounting and the Evolution of Selfing in Arenaria Uniflora (caryophyllaceae). Evolution, 54, 1558–1565. doi: 10.1111/j.0014-3820.2000.tb00701.x

Fishman, L., & Wyatt, R. (1999). Pollinator-Mediated Competition, Reproductive Character Displacement, and the Evolution of Selfing in Arenaria uniflora (Caryophyllaceae). Evolution, 53, 1723–1733. doi: 10.2307/2640435

Gervasi, D. D. L., & Schiestl, F. P. (2017). Real-time divergent evolution in plants driven by pollinators. Nature Communications, 8, 1–8. doi: 10.1038/ncomms14691

Goodwillie, C., Kalisz, S., & Eckert, C. G. (2005). The Evolutionary Enigma of Mixed Mating Systems in Plants: Occurrence, Theoretical Explanations, and Empirical Evidence. Annual Review of Ecology, Evolution, and Systematics, 36, 47–79.

Goodwillie, C., & Ness, J. M. (2013). Interactions of hybridization and mating systems: A case study in Leptosiphon (Polemoniaceae). American Journal of Botany, 100, 1002–1013. doi: 10.3732/ajb.1200616

Gröning, J., & Hochkirch, A. (2008). Reproductive Interference Between Animal Species. The Quarterly Review of Biology, 83, 257–282. doi: 10.1086/590510

Harder, L. D., Cruzan, M. B., & Thomson, J. D. (1993). Unilateral incompatibility and the effects of interspecific pollination for Erythronium americanum and Erythronium albidum (Liliaceae). Canadian Journal of Botany, 71, 353–358. doi: 10.1139/b93-038

Huang, S.-Q., & Shi, X.-Q. (2013). Floral isolation in Pedicularis: how do congeners with shared pollinators minimize reproductive interference? New Phytologist, 199, 858–865. doi: 10.1111/nph.12327

Hubbell, S. P. (2001). The Unified Neutral Theory of Biodiversity and Biogeography. Princeton University Press, Princeton.

Husband, B. C., & Schemske, D. W. (1996). Evolution of the Magnitude and Timing of Inbreeding Depression in Plants. Evolution, 50, 54–70. doi: 10.2307/2410780

Kalisz, S., Randle, A., Chaiffetz, D., Faigeles, M., Butera, A., & Beight, C. (2012). Dichogamy correlates with outcrossing rate and defines the selfing syndrome in the mixed-mating genus Collinsia. Annals of Botany, 109, 571–582. doi: 10.1093/aob/mcr237

Kalisz, S., Vogler, D. W., & Hanley, K. M. (2004). Context-dependent autonomous self-fertilization yields reproductive assurance and mixed mating. Nature, 430, 884–887. doi: 10.1038/nature02776

Karron, J. D., Jackson, R. T., Thumser, N. N., & Schlicht, S. L. (1997). Outcrossing rates of individual Mimulus ringens genets are correlated with anther–stigma separation. Heredity, 79, 365–370. doi: 10.1038/hdy.1997.169

Katsuhara, K. R., & Ushimaru, A. (2019). Prior selfing can mitigate the negative effects of mutual reproductive interference between coexisting congeners. Functional Ecology, 33, 1504–1513. doi: 10.1111/1365-2435.13344

Kishi, S., & Nakazawa, T. (2013). Analysis of species coexistence co-mediated by resource competition and reproductive interference. Population Ecology, 55, 305–313. doi: 10.1007/s10144-013-0369-2

Larson, B. M. H., & Barrett, S. C. H. (2000). A comparative analysis of pollen limitation in flowering plants. Biological Journal of the Linnean Society, 69, 503–520. doi: 10.1111/j.1095-8312.2000.tb01221.x

Levin, D. A., & Anderson, W. W. (1970). Competition for Pollinators between Simultaneously Flowering Species. The American Naturalist, 104, 455–467.

Lloyd, D. G. (1992). Self- and Cross-Fertilization in Plants. II. The Selection of Self-Fertilization. International Journal of Plant Sciences, 153, 370–380.

Lloyd, D. G., & Webb, C. J. (1986). The avoidance of interference between the presentation of pollen and stigmas in angiosperms I. Dichogamy. New Zealand Journal of Botany, 24, 135–162. doi: 10.1080/0028825X.1986.10409725

Martin, N. H., & Willis, J. H. (2007). Ecological Divergence Associated with Mating System Causes Nearly Complete Reproductive Isolation Between Sympatric Mimulus Species. Evolution, 61, 68–82. doi: 10.1111/j.1558-5646.2007.00006.x

May, R. M. (1974). Stability and complexity in model ecosystems. Princeton University Press, Princeton.

Mitchell, R. J., Flanagan, R. J., Brown, B. J., Waser, N. M., & Karron, J. D. (2009). New frontiers in competition for pollination. Annals of Botany, 103, 1403–1413. doi: 10.1093/aob/mcp062

Moreira-Hernández, J. I., & Muchhala, N. (2019). Importance of Pollinator-Mediated Interspecific Pollen Transfer for Angiosperm Evolution. Annual Review of Ecology, Evolution, and Systematics, 50, 191–217. doi: 10.1146/annurev-ecolsys-110218-024804

Nishida, S., Kanaoka, M. M., Hashimoto, K., Takakura, K.-I., & Nishida, T. (2014). Pollen–pistil interactions in reproductive interference: comparisons of heterospecific pollen tube growth from alien species between two native Taraxacum species. Functional Ecology, 28, 450–457. doi: 10.1111/1365-2435.12165

Randle, A. M., Spigler, R. B., & Kalisz, S. (2018). Shifts to earlier selfing in sympatry may reduce costs of pollinator sharing. Evolution, 72, 1587–1599. doi: 10.1111/evo.13522

Roels, S. A. B., & Kelly, J. K. (2011). Rapid Evolution Caused by Pollinator Loss in Mimulus Guttatus. Evolution, 65, 2541–2552. doi: 10.1111/j.1558-5646.2011.01326.x

Runquist, R. D. B., & Stanton, M. L. (2013). Asymmetric and frequency-dependent pollinator-mediated interactions may influence competitive displacement in two vernal pool plants. Ecology Letters, 16, 183–190. doi: 10.1111/ele.12026

Runquist, R. D. B. (2012). Pollinator-mediated competition between two congeners, Limnanthes douglasii subsp. rosea and L. alba (Limnanthaceae). American Journal of Botany, 99, 1125–1132. doi: 10.3732/ajb.1100588

Schemske, D. W., & Lande, R. (1985). The Evolution of Self-Fertilization and Inbreeding Depression in Plants. Ii. Empirical Observations. Evolution, 39, 41–52. doi: 10.1111/j.1558-5646.1985.tb04078.x

Sicard, A., & Lenhard, M. (2011). The selfing syndrome: a model for studying the genetic and evolutionary basis of morphological adaptation in plants. Annals of Botany, 107, 1433–1443. doi: 10.1093/aob/mcr023

Silvertown, J. (2004). Plant coexistence and the niche. Trends in Ecology & Evolution, 19, 605–611. doi: 10.1016/j.tree.2004.09.003

Takakura, K.-I., Nishida, T., Matsumoto, T., & Nishida, S. (2008). Alien dandelion reduces the seed-set of a native congener through frequency-dependent and one-sided effects. Biological Invasions, 11, 973–981. doi: 10.1007/s10530-008-9309-z

Tokuda, N., Hattori, M., Abe, K., Shinohara, Y., Nagano, Y., & Itino, T. (2015). Demonstration of pollinator-mediated competition between two native Impatiens species, Impatiens noli-tangere and I. textori (Balsaminaceae). Ecology and Evolution, 5, 1271–1277. doi: 10.1002/ece3.1431

van der Niet, T., & Johnson, S. D. (2012). Phylogenetic evidence for pollinator-driven diversification of angiosperms. Trends in Ecology & Evolution, 27, 353–361. doi: 10.1016/j.tree.2012.02.002

Webb, C. J., & Lloyd, D. G. (1986). The avoidance of interference between the presentation of pollen and stigmas in angiosperms II. Herkogamy. New Zealand Journal of Botany, 24, 163–178. doi: 10.1080/0028825X.1986.10409726

Whitton, J., Sears, C. J., & Maddison, W. P. (2017). Co-occurrence of related asexual, but not sexual, lineages suggests that reproductive interference limits coexistence. Proc. R. Soc. B, 284, 20171579. doi: 10.1098/rspb.2017.1579

